# Red squirrels mitigate costs of territory defence through social plasticity

**DOI:** 10.1101/453993

**Authors:** Erin R. Siracusa, David R. Wilson, Emily K. Studd, Stan Boutin, Murray M. Humphries, Ben Dantzer, Jeffrey E. Lane, Andrew G. McAdam

## Abstract

For territorial species, the ability to be behaviourally plastic in response to changes in their social environment may be beneficial by allowing individuals to mitigate conflict with conspecifics and reduce the costs of territoriality. Here we investigated whether North American red squirrels (*Tamiasciurus hudsonicus*) were able to minimize costs of territory defence by adjusting behaviour in response to the familiarity of neighbouring conspecifics. Since red squirrels living in familiar neighbourhoods face reduced intrusion risk, we predicted that increasing familiarity among territorial neighbours would allow squirrels to spend less time on territorial defence and more time in the nest. Long-term behavioural data (1995-2004) collected from the same squirrels across several different social environments indicated that red squirrels reduced rates of territorial vocalizations and increased nest use in response to increasing familiarity with neighbours. In contrast, cross-sectional data (2015-2016), which provided observations from each individual in a single social environment, did not provide evidence of this plasticity. Post-hoc analyses revealed that evidence of social plasticity in this system was primarily due to within-individual changes in behaviour, which we were unable to estimate in the cross-sectional data. Our results demonstrate that red squirrels can reduce the costs of territoriality by appropriately adjusting behaviour in response to changes in their social environment. However, our results also suggest that estimating plasticity by comparing behaviour among individuals (i.e. cross-sectional analyses) may not always be reliable. Our ability to detect these effects may therefore depend on having data with multiple observations from the same individuals across different social environments.

## Introduction

Phenotypic plasticity is one way for organisms to adjust their physiology, morphology, and behaviour when the rate of environmental change outpaces the rate of genetic evolutionary change (Piersma & Drent, 2003). Broadly defined, phenotypic plasticity is the ability of a given genotype to express multiple phenotypes in response to different environmental conditions (Pigliucci, 2001). Although classic studies of phenotypic plasticity have focused on changes in non-reversible traits (e.g. morphology) that are expressed within a single genotype (Hebert & Grewe, 1985; Lively, 1986; Greene, 1989), traits that are expressed repeatedly over the course of an organism’s lifetime (e.g. timing of reproduction) can be subject to reversible within-individual plasticity (Nussey, Wilson, & Brommer, 2007; Piersma & Drent, 2003). This ‘reversible plasticity’ (Gabriel, Luttbeg, Sih, & Tollrian, 2005), also referred to as ‘phenotypic flexibility’ (Piersma & Drent, 2003), or ‘responsiveness’ (Wolf, Van Doorn, & Weissing, 2008), is a powerful mechanism for adapting to changing and unpredictable environmental conditions. Behavioural traits, in particular, show capacity for substantial phenotypic lability in response to changing environmental conditions within an organism’s lifetime and can facilitate an organism’s ability to cope with both predictable and unpredictable variation in the environment (Ghalambor, Angeloni, & Carroll, 2010).

The social environment is arguably one of the most dynamic and variable realms of an individual’s environment, since high levels of unpredictability are inherent when interacting with other agents that can also exhibit plasticity in behaviour. Examples of social plasticity (changes in behaviour in response to changing social conditions; Sih, Chang, & Wey, 2014;1 Montiglio, Wey, Chang, Fogarty, & Sih, 2017) are prevalent. For instance, individuals adjust their level of aggression according to the perceived level of threat imposed by neighbours versus strangers (Temeles, 1994). Interacting individuals change their signaling behaviour in response to bystanders (‘audience effect’-Doutrelant, McGregor, & Oliveira, 2001; Pinto, Oates, Grutter, & Bshary, 2011). Behaviour may also be affected by previous social experiences such as ‘winner-loser effects’ (Hsu, Earley, & Wolf, 2006; Rutte, Taborsky, & Brinkhof, 2006), as well as by ‘eavesdropping’, in which bystanders extract information from interacting individuals (Earley, 2010; Mennill, Ratcliffe, & Boag, 2002; Oliveira, McGregor, & Latruffe, 1998).

The ability to adjust behaviour in response to social context should allow individuals to avoid costly interactions while appropriately engaging in other social interactions that might enhance fitness (Taborsky & Oliveira, 2012). This ability to show adaptive adjustments in social behaviour has been termed ‘social skill’ (Sih & Bell, 2008) or ‘social competence’ (Taborsky & Oliveira, 2012), although few studies have directly demonstrated fitness benefits of social plasticity (Han & Brooks, 2015; Patricelli, Uy, Walsh, & Borgia, 2002; Montiglio et al., 2017). Given the substantial number of social interactions that group-living species must navigate, the benefits of social plasticity are expected to be high in such species (Taborsky, Arnold, Junker, & Tschopp, 2012). However, solitary, territorial species may also benefit from appropriate adjustments in social behaviour, as being socially plastic may allow individuals to mitigate conflict with conspecifics and reduce the costs of territoriality. For example, gladiator frogs (*Hypsiboas rosenbergi*) adjust the timing of vocalizations in response to changing levels of conspecific competition. By reducing calling rates in response to changing social conditions, individuals can minimize an energetically costly behaviour (Höbel, 2015).

Solitary, territorial species, like their social counterparts, face variation in their social environments through their interactions with territorial neighbours. A well-described example of this variation is differences in familiarity with neighbours (Bebbington et al., 2017; Beletsky & Orians, 1989; Eason & Hannon, 1994; Grabowska-Zhang, Wilkin, & Sheldon, 2012). Long-term social relationships with neighbours have been presumed to be advantageous by minimizing renegotiation of territory boundaries and therefore reducing aggression as well as time and energy spent on territory defence (‘dear-enemy effect’; Fisher, 1954). However, most evidence in support of this cooperative phenomenon comes from experimental studies where individuals are exposed to a familiar and unfamiliar stimulus and a behavioural response is recorded (Temeles, 1994). We know less about how behavioural time budgets are affected by long-term social relationships under natural conditions when individuals may have to navigate territorial dynamics with multiple neighbours (but see Bebbington et al., 2017; Eason & Hannon, 1994).

In this study, we used a longitudinal dataset spanning eight years and cross-sectional data from two years to examine whether territorial North American red squirrels (*Tamiasciurus hudsonicus*, hereafter ‘red squirrels’) adjust their behaviour in response to their familiarity with their local social environment. Our long-term dataset contained multiple observations of the same individuals across different social environments, while our cross-sectional data represented an intensive snapshot of a large number of individuals at a single point in time (i.e. a single social environment for each individual), allowing us to compare across individuals to assess differences in behaviour. Although behavioural plasticity is fundamentally a within-individual phenomenon, it can be approximated by comparing among individuals in different environments (Legagneux & Ducatez, 2013; Slabbekoorn & Peet, 2003). While this among-individual approach is a useful tool (particularly where it is challenging or time-consuming to collect data on many individuals over several environments) it relies on the critical assumption that the among-individual relationship is an accurate representation of within-individual changes in behaviour.

Red squirrels are territorial rodents that defend year-round exclusive territories (Smith, 1968). In the Yukon, red squirrels cache white spruce cones (*Picea glauca*) in a larder hoard called a ‘midden’ at the center of their territory (Fletcher et al., 2010). This food cache is important for overwinter survival (Kemp & Keith, 1970; LaMontagne et al., 2013) and both sexes heavily defend these resources from conspecifics, primarily through territorial vocalizations called ‘rattles’ (Smith, 1978). Rattles function to deter intruders (Siracusa, Morandini, et al., 2017b) but are also individually unique (Digweed, Rendall, & Imbeau, 2012; Wilson et al., 2015). Rattles therefore carry important information about the local social environment, such as the identity or density of neighbouring conspecifics. Squirrels have been shown to use this acoustic information to increase rattling rates and vigilance and decrease nest use in response to increasing local density (Dantzer, Boutin, Humphries, & McAdam, 2012). Additionally, familiarity (length of time as neighbours) with territorial neighbours has been shown in this system to have direct effects on territory intrusion risk. Specifically, individuals living in neighbourhoods with higher average familiarity faced reduced intrusion risk (Siracusa, Boutin, et al., 2017a), consistent with the dear enemy phenomenon (Fisher, 1954).

Given these previous findings, we predicted that increasing familiarity with territorial neighbours would allow for decreased time spent on territorial defence as evidenced by (1) decreasing rattling rates and (2) reducing time spent vigilant for conspecifics. We also predicted that, as a squirrel’s familiarity with its neighbours increased, squirrels would increase the proportion of time spent in nest, as a proxy for time spent on offspring-care or self-maintenance. Changes in behaviour, as predicted above, would allow individuals to minimize aggression and reduce allocation of time and energy to territory defence under social conditions associated with reduced risk of territorial intrusion, and thus would be indicative of social competence in this species.

## Methods

We studied a natural population of North American red squirrels located in the southwest Yukon near Kluane National Park (61° N, 138° W) that has been monitored continuously since 1987 as part of the Kluane Red Squirrel Project (KRSP; McAdam, Boutin, Sykes, & Humphries, 2007). To assess social plasticity in red squirrels, we measured behaviour of individuals on three study grids characterized by open boreal forest where white spruce is the dominant tree species (Krebs et al. 2001).

Using the longitudinal dataset, we analyzed long-term focal animal observations (Altmann, 1974) of 41 red squirrels across eight years (between 1995 and 2004), collected on one unmanipulated control grid (Sulphur: SU; 40 ha). We also examined a cross-section of data by analyzing focal observations of 108 squirrels in one year (2016) on two unmanipulated control grids (Kloo: KL and SU; 40 ha each) and one food supplemented grid (Agnes: AG; 45 ha; see Dantzer et al., 2012 for a description of the food supplementation experiment). In addition to using focal animal observations, we measured the behaviour of squirrels by deploying accelerometers in 2016 and audio recorders in 2015 and 2016. On average, squirrels rattle once every 8.24 min (Dantzer et al., 2012), which makes capturing fine-scale adjustments in behaviour challenging, even with the intensive use of focal observations. Therefore, our intention was to use data from audio recorders and accelerometers as additional measures of social plasticity that might better capture fine-scale adjustments in behaviours such as rattling rates and nest use. This research was approved by the University of Guelph Animal Care Committee (AUP number 1807) and is in compliance with the ASAB/ABS *Guidelines for the Use of Animals in Research*. Details on these approaches are provided below.

### Measuring familiarity

In each year, we enumerated all squirrels living on our study areas and monitored individuals from March until August. We used a combination of live-trapping procedures and behavioural observations to track reproduction, identify territory ownership, and determine offspring recruitment from the previous year (see Berteaux & Boutin, 2000; McAdam et al., 2007 for a complete description of core project protocols). All study grids were staked and flagged at 30 m intervals, which allowed us to record the spatial location of all squirrel territories to the nearest 3 m. In this study system, territory locations were denoted based on the location of an individual’s midden, which approximates the center of a squirrel’s territory. We did not explicitly map territory boundaries for all individuals.

We trapped squirrels using Tomahawk live traps (Tomahawk Live Trap Co., Tomahawk, Wisconsin, USA) baited with all natural peanut butter. If previously tagged, the identities of the squirrels were determined from their unique alphanumeric metal ear tags (Monel #1; National Band and Tag, Newport, KY, USA), which they received in their natal nest at around 25 days of age. During the first capture of the season, we marked each squirrel by threading coloured wires through each ear tag, which allowed for individual identification of squirrels during behavioural observations. We censused the population twice annually (in mid-May and mid-August) and determined territory ownership through a combination of consistent live-captures of the same individual at the same midden and behavioural observations of territorial ‘rattle’ vocalizations (Smith 1978).

For each territory owner we defined the social neighbourhood to be all conspecifics whose middens were within a 130 m radius of the owner’s midden. One hundred and thirty meters is the furthest distance that red squirrel rattles are known to carry (Smith, 1978) and is similar to the 150 m distance found by (Dantzer et al., 2012) to be most relevant for local density effects of the social environment. We measured pairwise familiarity between the territory owner and each neighbour as the number of days that both individuals were within the same acoustic neighbourhood (i.e. occupied their current neighbouring territories). We then averaged pairwise measures of familiarity between the focal individual and each of the neighbours to provide a measure of the average familiarity of each individual with its entire acoustic neighbourhood (Siracusa, Boutin, et al., 2017a).

### Focal observations

Red squirrels are an ideal species for behavioural studies because they are diurnal, easy to locate visually or through acoustic cues, and habituate readily to the presence of humans. As part of the KRSP, we have recorded the behaviour of red squirrels through focal sampling of radio-collared individuals (model PD-2C, 4 g, Holohil Systems Ltd., Ontario, Canada) since 1994, although the sampling protocol has varied slightly across this period. For this study, we chose to use a subset of long-term behavioural data where focal observations were collected in a consistent manner by instantaneous sampling at 30-sec intervals for ten continuous minutes (Altmann, 1974) on a single control grid (SU; *N* = 487 10-min sessions over 41 individuals). We excluded any focals where the squirrel was out of sight for more than half the observation session (*N* = 8). These behavioural observations were available for female squirrels in 1995 (*N* = 41), 1996 (*N* = 10), 1997 (*N* = 25), 1999 (*N* = 34), 2001 (*N* = 70), 2002 (*N* = 110), 2003 (*N* = 120), and 2004 (*N* = 77) and were recorded by 38 different observers. On average, data were available for two different social environments per individual (range 1-8). We considered any change in average familiarity during the population census to be a change in the social environment. Since the population was censused twice annually (in mid-May and mid-August) individuals could experience up to two different social environments per year.

Between 7 May 2016 and 31 August 2016, we used focal animal sampling as described above for seven continuous minutes, rather than ten, to record red squirrel behaviour (*N* = 1060 7 min sessions over 108 individuals). In this cross-sectional data we only had observations from each individual in a single social environment. Since rattling is a rare behaviour and is often missed using instantaneous sampling, in 2016 we recorded all occurrences (Altmann, 1974) of rattle vocalizations emitted by the focal squirrel, including those which fell outside the 30-sec sampling interval (i.e ‘critical incidents’). We used all of these data, including critical incidents, to assess how familiarity affected rattling rates in 2016. Four observers collected behavioural data on both male (*N* = 76) and female (*N* = 32) squirrels across two control grids (KL and SU) and one food-supplemented grid (AG). We monitored each individual for 2-10 days consecutively (barring inclement weather; mean = 4 days) and collected an average of 10 focals per individual (range: 2-29). In instances where multiple focal observations were collected for the same squirrel in a single day, observations were kept 30 minutes apart at minimum. Because an observer was in regular attendance at these territories we could be confident that there was no turnover in the social environment during the sampling period for any of these individuals. Territory turnovers in this system are accompanied by substantial rattling and chasing and are therefore easy to detect. The two squirrels for which we observed a disturbance in the local social environment during the sampling period were excluded from this analysis.

For all focal sampling, we recorded and classified red squirrel behaviours in a similar way to previous studies of squirrel behaviour in this system (Anderson & Boutin, 2002; Dantzer et al., 2012; Stuart-Smith & Boutin, 1994). We classified behaviours according to the following categories: vocalizing (“barking” or “rattling”; Smith, 1978), feeding, foraging, traveling, caching food items, interacting with conspecifics, grooming, resting, vigilant, in nest, or out of sight (unknown behaviour). Vigilance could be distinguished from resting by the alert posture of the squirrel; vigilant squirrels typically had their head up and appeared observant, sometimes standing on their hind limbs, while resting squirrels often had their head tucked down or lay stretched out.

### Audio recording and acoustic analysis

Between 23 June – 25 September 2015 and 8 May – 1 September 2016, we deployed Zoom H2n audio recorders (Zoom Corporation, Tokyo, Japan) to determine rattling rates of squirrels. We attached recorders with windscreens to 1.5 m stakes and placed a single recorder in the center of each squirrel’s midden. Since Zoom H2n recorders are not weatherproof, we placed an umbrella approximately 30 cm above each audio recorder to protect it from rain and snow. Each morning, we deployed audio recorders between 0500-0600 h (just before squirrels typically became active). We set audio recorders to record in 44.1kHz/16bit WAVE format, and recorded in 2-channel surround mode with the stereo width set to +6 dB and the mic gain set to 10. We allowed audio recorders to run for a full 24 hours, but in this study we only use data collected between 0700-1300 h, which is the period during which squirrels are typically most active between early summer and early autumn (Studd, Boutin, McAdam, & Humphries, 2016; Williams et al., 2014). We deployed audio recorders for 137 squirrels (*N* = 109 males and *N* = 28 females) and recorded each squirrel for 5 consecutive days on average (range: 1-13 days; *N* = 714 days or 4284 hours over 137 individuals). Because we collected audio data over 2 years, we had observations from 2-3 different social environments for 28 of these individuals. Due to the large volume of recordings, we detected rattle vocalizations from recordings automatically using Kaleidoscope software (version 4.3.2; Wildlife Acoustics, Inc., Maynard, MA, USA). Detection settings included a frequency range of 2000–13000 Hz, a signal duration of 0.4–15 s, a maximum intersyllable silence of 0.5 s, a fast Fourier transform size of 512 points (corresponding to a frequency resolution of 86 Hz and a temporal resolution of 6.33 ms), and a distance setting of 2 (this value ensures that all detections are retained).

The purpose of using audio recorders was to provide a more accurate estimate of individual rattling rates. A challenge, however, is that the recorders also recorded vocalizations from neighbouring squirrels. Neighbours should be farther away from the recorder. Because sound degrades and attenuates predictably with distance, it should be possible to distinguish between the rattles of focal and neighbour squirrels on the basis of rattle acoustic structure. We tested this by conducting hour-long calibrations on 48 focal individuals between 13 September and 14 October 2015. During these calibrations, audio recorders were set up as described above. A single observer standing near the midden kept the territory owner in sight and recorded whether each rattle belonged to the territory owner or a neighbouring individual.

We detected rattle vocalizations from the calibration recordings using Kaleidoscope software (same settings as above). Based on a comparison with the observer’s notes, the software detected 100% of the focal squirrel rattles. We then developed a procedure for distinguishing focal squirrel rattles from other types of detections, including neighbour rattles and non-rattles. First, we automatically measured the acoustic structure of every detection using the software package ’Seewave’ (version 2.0.5; Sueur, Aubin, & Simonis, 2008) in R (see details of structural measures below). Second, we used the structural measurements in a discriminant function analysis in SPSS (software, version 24; IBM Corporation, Armonk, New York, USA) to develop a predictive model for assigning detections to groups (*i.e.,* focal rattle, neighbour rattle, non-rattle). We developed the model using detections from half of the 1-hour calibration files (selected at random), and then tested it for accuracy by applying it to the detections from the remaining half. The model correctly assigned 80.6% of the focal rattles to the ’focal rattle’ group, meaning we missed 19.4% of focal rattles (i.e. false negatives). Some non-rattle detections were also assigned to the ’focal rattle’ group, but we removed these by reviewing the spectrograms of all detections categorized as ‘focal rattles’. After removing non-rattle detections, 16.0% of the detections remaining in the ’focal rattle’ group were false positives, meaning they were actually from the neighbour instead of the focal squirrel. We then applied the predictive model to the main set of audio files, and reviewed all detections labeled as ‘focal rattle’ in Kaleidoscope to remove the non-rattle detections.

The structural measures included in the discriminant function analysis were: (1) duration, (2) root-mean-square amplitude, (3) pulse rate, (4) duty cycle, and five variables that measured the distribution of energy in the frequency domain, including (5) peak frequency, (6) first energy quartile, (7) skewness, (8) centroid, and (9) and spectral flatness. Duration, root-mean-square amplitude, pulse rate, and duty cycle were measured from a waveform. Pulse rate is the number of pulses in the rattle minus one, divided by the period of time between the beginning of the first pulse and the beginning of the last (as in Wilson et al., 2015). Duty cycle is the proportion of the rattle when a pulse is being produced. For pulse rate and duty cycle, individual pulses were identified using the ’timer’ function in seewave (50% amplitude threshold; 200-point smoothing window with 90% overlap). The five energy distribution variables were obtained using the ’specprop’ function in seewave, and were based on a mean frequency spectrum (512-point fast Fourier transform, hanning window, 0% overlap). Peak frequency is the frequency of maximum amplitude. First energy quartile is the frequency below which 25% of the energy is found. Skewness, centroid, and kurtosis describe the shape of the power spectrum (detailed definitions can be found in Sueur et al., 2008).

### Accelerometers

Between 4 May and 1 September 2016, we deployed AXY-3 accelerometers (Technosmart Europe srl., Rome, Italy) on 94 squirrels (*N* = 66 males and *N* = 28 females). An accelerometer is an instrument that measures the acceleration of the body along three axes: anterior-posterior (surge), lateral (sway), dorso-ventral (heave) and records temperature, allowing for the characterization of different behavioural patterns. Accelerometers were deployed in combination with radio transmitters (model PD-2C, Holohil Systems Ltd., Ontario, Canada). Total package weight for collars with both accelerometers and radio transmitters (including battery, packaging, and bonding material) was 9.6 g on average. For a 200-250 g red squirrel (Steele, 1998) this collar weight was less than the recommended 5-10% of the animal’s body weight (Wilson, Cole, Nichols, Rudran, & Foster, 1996). We deployed accelerometers on 94 individuals for an average of 9 days per individual (range: 4-17; *N* = 873 days over 94 individuals) at a sampling rate of 1 Hz. Accelerometers recorded data constantly while deployed, but for this study we only use data between 06:00-21:00 h to estimate time spent in nest during active hours of the day (Williams et al., 2014).

Raw accelerometer data were classified into 5 behavioural categories using threshold values of summary statistics according to the decision tree developed for red squirrel accelerometers and temperature data loggers by Studd et al. (in review). Following methods proposed by Collins et al. (2015), the decision tree was created using 83.8 hours of direct behavioural observations on 67 free-ranging squirrels and had an overall accuracy of correctly classifying known behaviours of 94.9% (Studd et al. in review). Briefly, warm stable temperatures were used to identify when the animal was in the nest with the additional constraint that the individual must not be moving for the majority of each nest bout. Low acceleration values were associated with not moving, moderate acceleration values denoted feeding, and high acceleration corresponded to travelling. Travelling was further categorized as running when the peak acceleration value of the surge axis was above a threshold of 1.15 G.

### Statistical analyses

Given that previous work in this study system (Dantzer et al., 2012) allowed us to make specific predictions about how squirrels should adjust rattling rates, vigilance and nest use in response to their social environment, here we used univariate models to test for the effects of familiarity on each of these behaviours explicitly. For all models we included local density, measured as the number of squirrels per hectare within 130 m, as a continuous predictor, to account for the fact that previous work in the study system has found local density to be an important predictor of behavioural time budgets (Dantzer et al., 2012). We also included age as a fixed effect in all rattling rate models since we expected that the vigor of territory defence might decline with physical deterioration, but we did not have specific predictions as to how age might affect nest use or vigilance. However, it is important to note that since young squirrels are inherently unfamiliar with their neighbours and familiarity increases with age, age and familiarity were strongly correlated (Pearson’s correlation coefficient ranged between 0.42 and 0.58 for these analyses) although variance inflation factors were low (< 3; Zuur, Ieno, & Elphick, 2009). Fixed effects and random effects of all models are summarized in Tables 1 and 2, respectively.

#### Focal data

To account for the structural differences in our data sets, multiple observations of the same individuals across different social environments (longitudinal data) versus observations from different individuals, each in a single social environment (cross-sectional data), we analyzed the longitudinal (*N* = 487 10-min sessions) and cross-sectional (*N* = 1060 7-min sessions) focal data separately. In the long-term data, there was a single data point where the number of rattles recorded was 25 times greater than the mean. This outlier was likely an error in data entry and was removed (see Figure S1). We analyzed the effects of neighbourhood familiarity on (1) the frequency of territorial vocalizations (rattles), (2) the proportion of time spent vigilant, and (3) the proportion of time spent in nest. We modeled the frequency of territorial vocalizations using a generalized linear mixed-effect model (GLMM) with a bobyqa optimizer and a Poisson error distribution (log-link) where the response variable was the number of rattles emitted during the 10-min focal session. For both the proportion of time spent in nest and the proportion of time spent vigilant, we fitted a Beta-Binomial model to account for overdispersion in the data (Harrison, 2015). Using the ‘cbind’ function, we defined the response variable as a 2-column matrix composed of the number of observations of the given behaviour (in nest or vigilant) and the number of observations of all other behaviours (not including observations when the squirrel was out of sight). In all models we included average familiarity and local density as continuous predictors, and for the rattling rate models we included age as a continuous fixed effect. We included grid, sex and observer identity as categorical fixed effects for the 2016 focal data (it was not necessary to include grid or sex for the long-term data as all data were collected on females on a single grid). For both datasets, we included a random intercept term for squirrel identity (squirrel ID) to account for repeated observations of the same squirrels. We also included a random effect of year and observer identity for the long-term dataset to account for inter-individual differences in behavioural scoring.

#### Audio recorder data

To assess the effects of familiarity on rattling rates derived from the audio recorder data, we fitted a GLMM with a Poisson error distribution (log-link). Our response variable was the number of rattles emitted between 0700 – 13:00 h (i.e. number of ‘focal rattles’, unadjusted for false positive or false negative error rates; *N* = 714 days of recordings). We included average familiarity, local density, age, grid, and sex as covariates in the model, as well as a random intercept term for squirrel ID, and an observation-level random effect (OLRE) to account for overdispersion in the model.

#### Accelerometer data

Using accelerometer data, we assessed the effect of neighbourhood familiarity on the proportion of time spent in nest between 06:00 – 21:00 h using a Beta-Binomial model (*N* = 873 days). Our response variable was defined as above, using a two-column matrix that included the number of nest observations and the number of observations of all other behaviours. We included average familiarity, local density, grid, and sex as fixed effects in the model, and included a random effect for squirrel ID and accelerometer collar.

#### Exploratory post-hoc analysis

Upon finding evidence of behavioural plasticity in the long-term data but not the cross-sectional data (see results below), we conducted an exploratory post-hoc analysis in an attempt to understand the inconsistencies in our results. While the longitudinal data provided multiple measures of the same individuals across different social environments, allowing us to estimate within-individual relationships, we could only estimate among-individual relationships in the cross-sectional data. To assess whether our results might be driven by a within-individual effect, thus limiting our ability to detect behavioural plasticity in the cross-sectional data, we re-fit our rattling rate and nest use models from the long-term data using a within-subject mean centering approach. Following the methodology of van de Pol & Wright (2009), we split our familiarity term into an among-individual effect of familiarity (i.e. the mean familiarity score for an individual across all observations) and a within-individual effect of familiarity (i.e. the deviation in each familiarity observation for each individual from their mean score). We applied the same approach to the 2015 and 2016 audio recorder data for which we had observations from individuals across multiple social environments (Table 3).

We conducted analyses using R version 3.4.1 (R Core Team, 2017) and fitted all GLMMs using the lme4 package (version 1.1-13; Bates, Maechler, Bolker, & Walker, 2015). For all analyses, we fitted generalized additive models to confirm that there were no significant non-linearities between our predictor and response variables. We checked for overdispersion by comparing the ratio of the sum of the squared Pearson residuals to the residual degrees of freedom in each model (Zuur et al., 2009) and assessing whether the sum of squared Pearson residuals approximated a Chi-squared distribution with n-p degrees of freedom (Bolker et al., 2009). As stated above, we accounted for overdispersion in Poisson models by including an observation-level random effect (OLRE; Harrison, 2014). For models with binomial data, we accounted for overdispersion using Beta-Binomial models, which have been demonstrated to better cope with overdispersion in binomial data (Harrison, 2015). We fitted all Beta-Binomial models using the package glmmADMB (version 0.8.3.3; Harrison, 2015; Skaug, Fournier, Nielsen, Magnusson, & Bolker, 2018). We standardized all continuous fixed effects to a mean of zero and unit variance. For the following results we present all means ± SE, unless otherwise stated, and consider differences statistically significant at P < 0.05.

## Results

Among the years in which we analyzed long-term focal data (1995-2004), variation in average neighbourhood familiarity ranged from 0 (corresponding to when a squirrel first established its territory) to 813 days (mean: 229 ± 9 days) and variation in local density ranged from 0.57 to 5.84 squirrels/hectare (mean: 1.93 ± 0.05 squirrels/hectare). In our 2015 and 2016 data, there was a nearly equivalent amount of variation in average neighbourhood familiarity and local density. Neighbourhood familiarity ranged from 0 to 855 days (mean: 296 ± 5 days) and local density ranged from 1.13 to 6.03 (mean: 3.34 ± 0.03 squirrels/hectare). Below we discuss the effects of familiarity and age on behavioural patterns. Results for other fixed effects in the models can be found in Table 1.

### Longitudinal data

#### Territorial defence

During the long-term focal observations, red squirrels emitted an average of 0.37 ± 0.04 rattles per 10-min observation session (range: 0-4), which is equivalent to one rattle every 27.06 minutes. (Rattling rates were much lower than in the cross-sectional data (see below) due to differences in behavioural sampling protocol. In 2016, all occurrences of rattling were recorded as ‘critical incidents’, while in the long-term data rattles were only recorded if they fell on a 30-second sampling interval. When critical incidents of rattling were removed from the 2016 data, rattling rates dropped to one rattle every 40.55 minutes.) Red squirrels in the longitudinal dataset adjusted their behaviour in response to increasing average neighbourhood familiarity by emitting significantly fewer rattles (*β* = −0.29 ± 0.12, *z* = −2.48, *P* = 0.01; Figure 1). This corresponds to a predicted three-fold decrease in rattling rates: in neighbourhoods with the lowest familiarity, squirrels were predicted to rattle once every 24.76 minutes and in neighbourhoods with the highest familiarity, only once every 79.75 minutes. The effect of age on rattling rates was marginally non-significant (*β* = −0.20 ± 0.11, *z* = −1.85, *P* = 0.06; Table 1). On average, squirrels spent 6.0 ± 0.7% of their time vigilant, but did not show changes in vigilance behaviour in response to changing familiarity with neighbours (*β* = 0.02 ± 0.13, *z* = 0.15, *P* = 0.88; Table 1).

#### Nest use

Based on the long-term data, red squirrels spent, on average, 31.0 ± 2.0% of their time in nest. Red squirrels responded to changing social conditions by increasing nest use in response to increasing familiarity (*β* = 0.26 ± 0.12, *z* = 2.31, *P* = 0.02; Figure 1). This is equivalent to a predicted 24% increase in nest use: squirrels in neighbourhoods with the lowest familiarity were predicted to spend only 19% of their time in nest compared to 43% in neighbourhoods with the highest familiarity.

### Cross-sectional data

#### Territorial defence

During focal observations in 2016, red squirrels emitted 0.71 ± 0.03 rattles per 7-min observation session (range: 0-6), which equates to approximately one rattle every 9.80 minutes. Data from audio recorders in 2015 and 2016 provided very similar estimates of rattling rates. We captured, on average, 33.96 ± 0.72 rattles per 6-hours of recording (range: 3-123), which, after correcting for the error rates in our discriminant function analysis, is equivalent to one rattle every 9.81 minutes. Based on both cross-sectional focal observations and audio recorder data, neither average familiarity of the social neighbourhood (all |*z*| < 1.25, all *P* > 0.21) nor age (all |*z*| < 1.87, all *P* ≥ 0.06) were significant predictors of rattling rate (Table 1). Focal observations indicated that red squirrels spent 7.0% ± 0.5% of their time vigilant, on average, but did not adjust vigilance behaviour in response to changing familiarity with neighbours (*β* = 0.05 ± 0.07, *z* = 0.69, *P* = 0.49; Table 1).

#### Nest use

Based on focal observations in 2016, red squirrels spent an average of 36.0% ± 1.0% of their time in nest. Accelerometer data from 2016 provided similar estimates of average proportion of time spent in nest during daylight hours (36.0% ± 0.4%). Both focal observations and accelerometer data indicated that squirrels did not adjust their nest use in response to familiarity with neighbours (all |*z*| < 1.22, all *P* > 0.22; Table 1).

### Exploratory post-hoc analysis

In our post-hoc analyses we found evidence to suggest that effects of familiarity on rattling rates were primarily due to within-individual changes in behaviour rather than among-individual differences. In the long-term data, increasing familiarity led to a significant decrease in rattling rates within (*β* = −0.21 ± 0.08, *z* = −2.51, *P* = 0.01), but not among individuals (*β* = −0.18 ± 0.12, *z* = −1.50, *P* = 0.13; Table 3). There was a positive within and among-individual effect of familiarity on nest use, but neither of these effects were significant (all |*z*| < 1.69, all *P* > 0.08; Table 3). Audio recorder data from 2015 and 2016 also a revealed a significant negative within-individual effect (*β* = −0.03 ± 0.01, *z* = −2.55, *P* = 0.01), but not among-individual effect of familiarity on rattling rates (*β* = 0.02 ± 0.05, *z* = 0.34, *P* = 0.74; Table 3). Results from the audio data should be interpreted with caution as the inclusion of year in the model affected these results (see Table S1).

**Figure 1.**
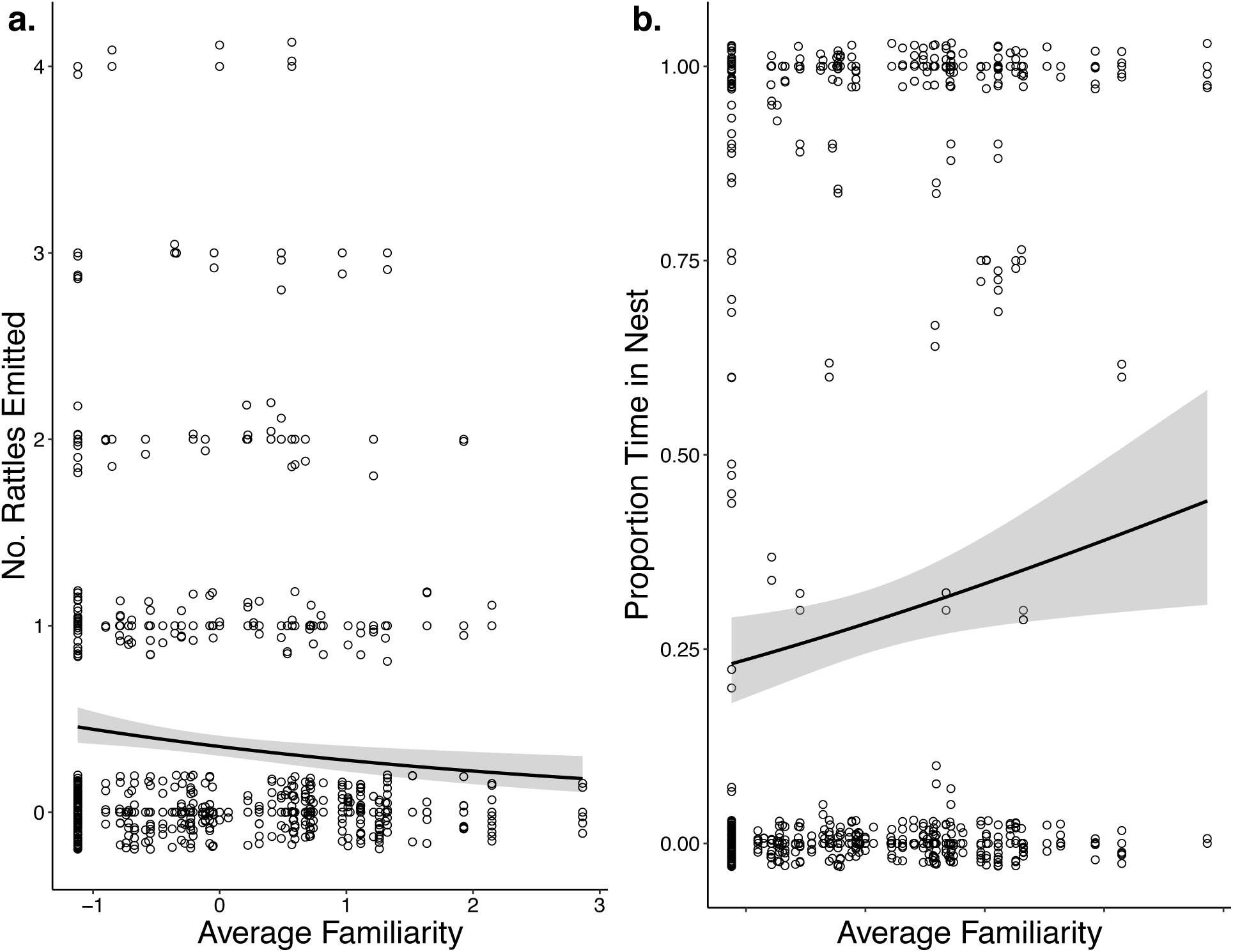
Red squirrels adjust a) rattling rate and b) proportion of time spent in nest in response to the average familiarity of their social neighbourhood (*N* = 487). Results are based on 10-min behavioural observations of squirrels between 1995-2004. Values on x-axis are standardized measures of average familiarity. Points indicate raw data with a small amount of jitter introduced to show overlapping points.

**Table 1.**
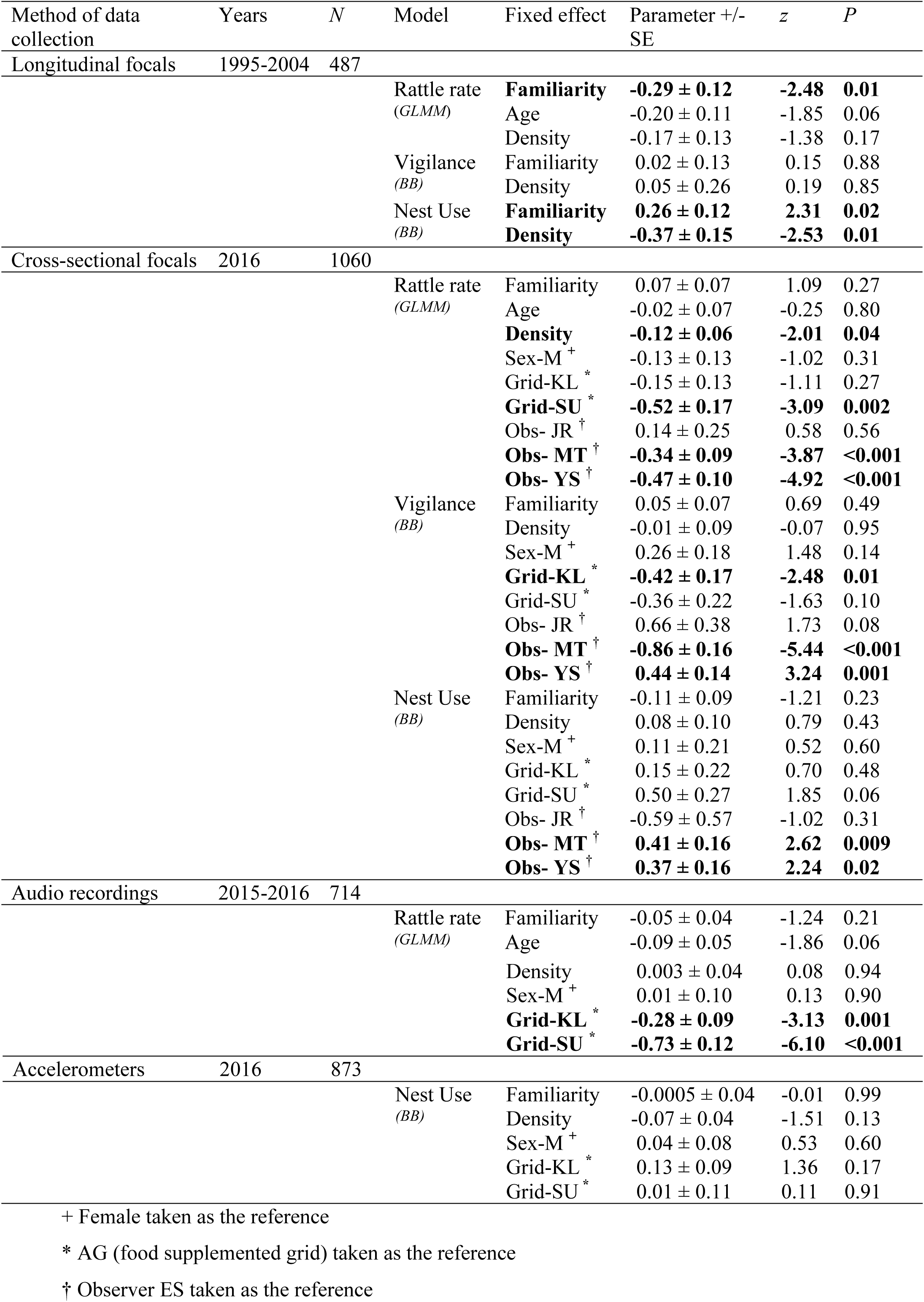
Fixed effects from all generalized linear mixed-effects (*GLMM*) and Beta-Binomial (*BB*) models, showing effects of average neighbourhood familiarity, local density, and focal squirrel’s age on rattling rate, nest use and vigilance behaviour. Regression coefficients for familiarity, age and density are standardized. Significant effects are indicated in bold.

**Table 2.**
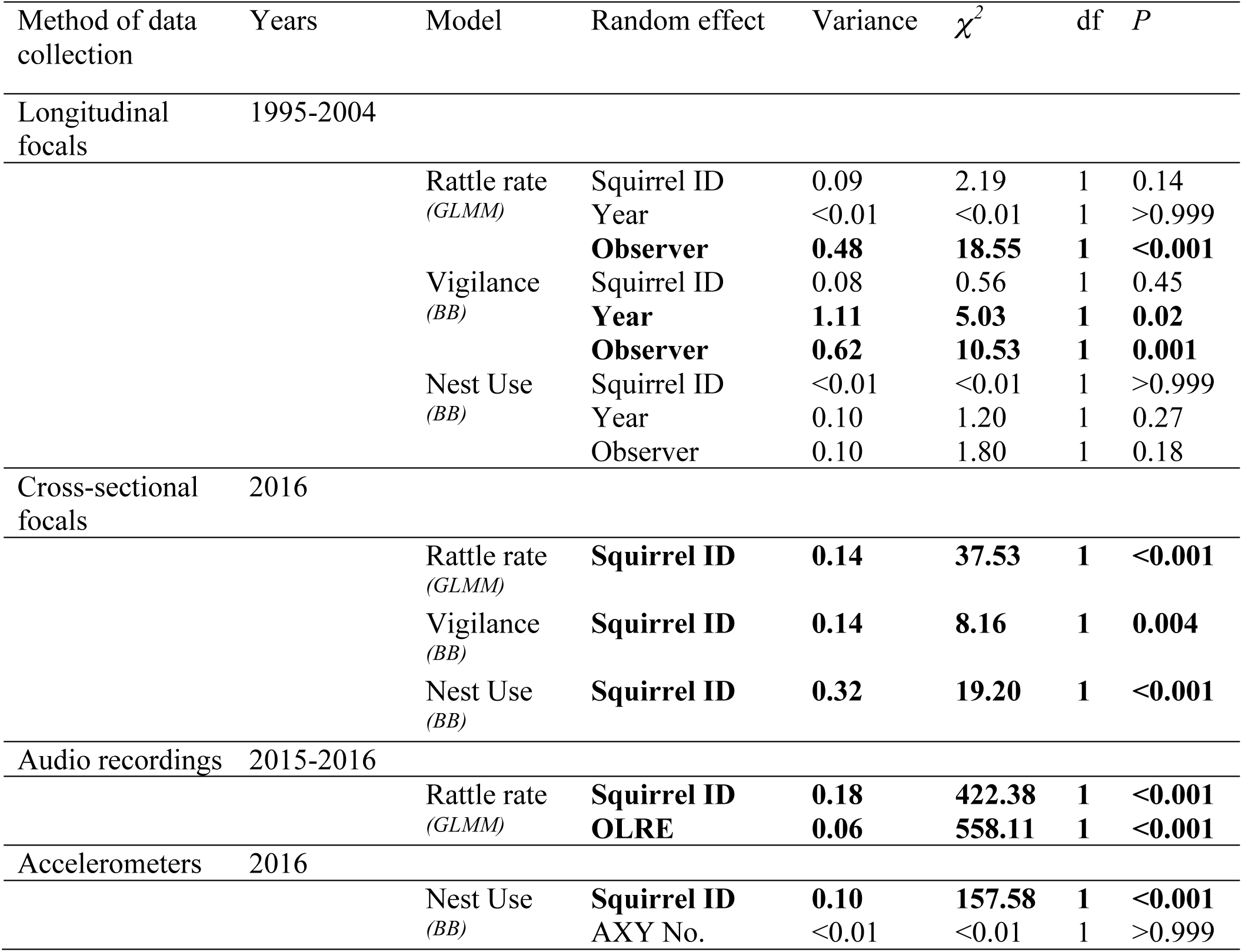
Random effects from all generalized linear mixed-effects (*GLMM*) and Beta-Binomial (*BB*) models. Significance assessed using a log-likelihood ratio test (LRT) with one degree of freedom to compare models with and without the listed random effect. Significant effects are indicated in bold.

| Method of data collection | Years | Model | Random effect | Variance | $\chi^2$ | df | *P* |
| --- | --- | --- | --- | --- | --- | --- | --- |
| Longitudinal focals | 1995-2004 |  |  |  |  |  |  |
|  |  | Rattle rate (*GLMM*) | Squirrel ID | 0.09 | 2.19 | 1 | 0.14 |
|  |  |  | Year | <0.01 | <0.01 | 1 | >0.999 |
|  |  |  | **Observer** | **0.48** | **18.55** | **1** | **<0.001** |
|  |  | Vigilance (*BB*) | Squirrel ID | 0.08 | 0.56 | 1 | 0.45 |
|  |  |  | **Year** | **1.11** | **5.03** | **1** | **0.02** |
|  |  |  | **Observer** | **0.62** | **10.53** | **1** | **0.001** |
|  |  | Nest Use (*BB*) | Squirrel ID | <0.01 | <0.01 | 1 | >0.999 |
|  |  |  | Year | 0.10 | 1.20 | 1 | 0.27 |
|  |  |  | Observer | 0.10 | 1.80 | 1 | 0.18 |
| Cross-sectional focals | 2016 |  |  |  |  |  |  |
|  |  | Rattle rate (*GLMM*) | **Squirrel ID** | **0.14** | **37.53** | **1** | **<0.001** |
|  |  | Vigilance (*BB*) | **Squirrel ID** | **0.14** | **8.16** | **1** | **0.004** |
|  |  | Nest Use (*BB*) | **Squirrel ID** | **0.32** | **19.20** | **1** | **<0.001** |
| Audio recordings | 2015-2016 |  |  |  |  |  |  |
|  |  | Rattle rate (*GLMM*) | **Squirrel ID** | **0.18** | **422.38** | **1** | **<0.001** |
|  |  |  | **OLRE** | **0.06** | **558.11** | **1** | **<0.001** |
| Accelerometers | 2016 |  |  |  |  |  |  |
|  |  | Nest Use (*BB*) | **Squirrel ID** | **0.10** | **157.58** | **1** | **<0.001** |
|  |  |  | AXY No. | <0.01 | <0.01 | 1 | >0.999 |

**Table 3.** Fixed effects from exploratory post-hoc models including a within-individual (Familiarity_*W*_) and among-individual (Familiarity_*A*_) effect of familiarity. Regression coefficients for familiarity, age and density are standardized. Significant effects are indicated in bold.

| Method of data collection | Years | *N* | Model | Fixed effect | Parameter<br>+/- SE | *z* | *P* |
| --- | --- | --- | --- | --- | --- | --- | --- |
| Longitudinal focals | 1995-2004 | 487 |  |  |  |  |  |
|  |  |  | Rattle rate<br>(GLMM) | **Familiarity<sub>W</sub>** | **-0.21 ± 0.08** | **-2.51** | **0.01** |
|  |  |  |  | Familiarity<sub>A</sub> | -0.18 ± 0.12 | -1.50 | 0.13 |
|  |  |  |  | **Age** | **-0.22 ± 0.11** | **-2.02** | **0.04** |
|  |  |  |  | Density | -0.18 ± 0.13 | -1.42 | 0.15 |
|  |  |  | Nest Use<br>(BB) | Familiarity<sub>W</sub> | 0.17 ± 0.10 | 1.68 | 0.09 |
|  |  |  |  | Familiarity<sub>A</sub> | 0.19 ± 0.13 | 1.53 | 0.13 |
|  |  |  |  | **Density** | **-0.37 ± 0.15** | **-2.50** | **0.01** |
| Audio recordings | 2015-2016 | 714 |  |  |  |  |  |
|  |  |  | Rattle rate<br>(GLMM) | **Familiarity<sub>W</sub>** | **-0.03 ± 0.01** | **-2.55** | **0.01** |
|  |  |  |  | Familiarity<sub>A</sub> | 0.02 ± 0.05 | 0.34 | 0.74 |
|  |  |  |  | **Age** | **-0.10 ± 0.05** | **-2.09** | **0.04** |
|  |  |  |  | Density | -0.02 ± 0.04 | -0.56 | 0.58 |
|  |  |  |  | Sex-M<sup>+</sup> | 0.01 ± 0.10 | 0.12 | 0.90 |
|  |  |  |  | **Grid-KL<sup>\*</sup>** | **-0.28 ± 0.09** | **-3.16** | **0.002** |
|  |  |  |  | **Grid-SU<sup>\*</sup>** | **-0.81 ± 0.12** | **-6.58** | **<0.001** |

+ Female taken as the reference
\* AG (food supplemented grid) taken as the reference

## Discussion

For territorial species, the ability to be responsive to changes in the social environment may convey a fitness advantage by allowing individuals to reduce time and energy investment in costly behaviours (Höbel, 2015; Krobath, Römer, & Hartbauer, 2017; Ydenberg, Giraldeau, & Falls, 1988). In this study, we used multiple types of behavioural data, as well as a longitudinal and cross-sectional dataset, to test a single overarching hypothesis: that red squirrels show behavioural plasticity in response to the familiarity of their social neighbourhood. Our results provide evidence that an ‘asocial’ species, the North American red squirrel, can respond to changes in the composition of its social environment, and that red squirrels do so under natural conditions and in a manner that is consistent with our expectations for adaptive behavioural change in this species.

Previous work in this study system has demonstrated that red squirrels face reduced intrusion risk in social neighbourhoods with high average familiarity (Siracusa, Boutin, et al., 2017a). As such, we predicted that red squirrels would show appropriate social plasticity by reducing territorial defence behaviours and increasing time and energy spent on self-maintenance behaviours when familiar with neighbouring conspecifics. Results from behavioural observations across eight years provided support for these predictions, indicating that red squirrels demonstrated social plasticity by reducing rattling rates and increasing the proportion of time spent in nest in social neighbourhoods with high average familiarity (Figure 1). Such changes in behaviour not only minimize the time spent on territory defence but might also reduce associated costs of territoriality. Territorial vocalizations may attract the attention of predators (Abbey-Lee, Kaiser, Mouchet, & Dingemanse, 2016) and rattles are loud, broadband signals which should be easy to localize (Marler 1955). By reducing rattling rates under less risky social conditions, squirrels may also benefit from reduced predation risk.

We did not, however, find effects of neighbourhood familiarity on vigilance behaviour. This could be due to vigilance for conspecifics being easily confounded with vigilance for predators. In contrast, Dantzer et al. (2012) found significant effects of local density on vigilance using behavioural data collected over a similar time frame, indicating that conspecific rather than heterospecific effects on vigilance are detectable in this system. While we included local density as a covariate in all of our models to account for the potential effects of density on behaviour (Dantzer et al., 2012), our goal was not to directly estimate effects of density, and our results therefore are not a clean representation of density effects. In several cases density was correlated with other variables in the model, such as grid, leading to substantial changes in the parameter estimates for density. As a result, the effects of density on behaviour that we have reported here cannot be compared directly to previous studies of these effects in this population (e.g. Dantzer et al., 2012; Shonfield, Taylor, Boutin, Humphries, & McAdam, 2012) and we do not discuss the effects of density further.

Results from the cross-sectional data in 2015 and 2016 did not corroborate our long-term results showing behavioural responses to familiarity. Findings from the behavioural observations, audio recorders, and accelerometers indicated that when using among-individual relationships to estimate the effects of the social environment on behaviour, there was no effect of familiarity on territorial behaviours (rattling rates, vigilance) or self-maintenance (nest use; Table 1). In light of this inconsistency with the longitudinal data, we conducted an exploratory post-hoc analysis to assess whether using longitudinal data (where we can estimate within-individual changes in behaviour) rather than cross-sectional data (where the analysis is largely among individuals) might account for our different findings. Since behavioural plasticity is functionally a within-individual phenomenon, using among-individual differences in behaviour to estimate these effects relies on the assumption that the among-individual relationship is an accurate representation of the within-individual relationship. Here we used a within-subject centering approach (van de Pol & Wright, 2009) and found that the within and among-individual effects were not equivalent. In the long-term data, we found that individuals adjusted rattling rates in response to changes within their own social environment (i.e. a significant within-individual effect) but did not observe significant differences in rattling rates when comparing among individuals (Table 3). Similarly, for the audio recorder data (the only cross-sectional data for which we had observations of individuals across multiple social environments) we found evidence of a significant within-individual, but not among-individual, effect (Table 3). Thus, while we clearly see evidence of plasticity when considering changes in individual behaviour across different social environments, in this study system it appears that we cannot estimate these effects by comparing behaviour among individuals.

One potential explanation for this discrepancy is that the among-individual effect is masked by individual variation in plasticity, whereby substantially different individual ‘slopes’ result in a ‘mean slope’ of zero (i.e. the absence of a significant population-level response to the environment; Nussey et al., 2007). We were unable to test this hypothesis as we lacked the statistical power to include a random slope term in our models (Martin, Nussey, Wilson, & Réale, 2011). Furthermore, even if all individuals demonstrate negative reaction norms (i.e. reduced rattling rate in response to increasing familiarity), there are still several reasons we might fail to detect differences among individuals. First, it seems unlikely that squirrels can assess their absolute familiarity, meaning that behavioural adjustments are dependent on the relative social environments individuals experience rather than absolute changes in familiarity. Additionally, variation in individual mean ratting rates (i.e. random intercepts) due to differences in sex, age, personality, stress, among other possibilities, might mask an among-individual effect. These factors, combined with variation in the range of social environments sampled for a given individual, mean that, even when all individuals show negative reaction norms, it is possible to measure a lack of (Figure 2b), or even a positive among-individual effect (Figure 2c). Additional individual data, spanning a range of social environments, is necessary to better understand the patterns leading to within versus among-individual effects in this system.

**Figure 2.**
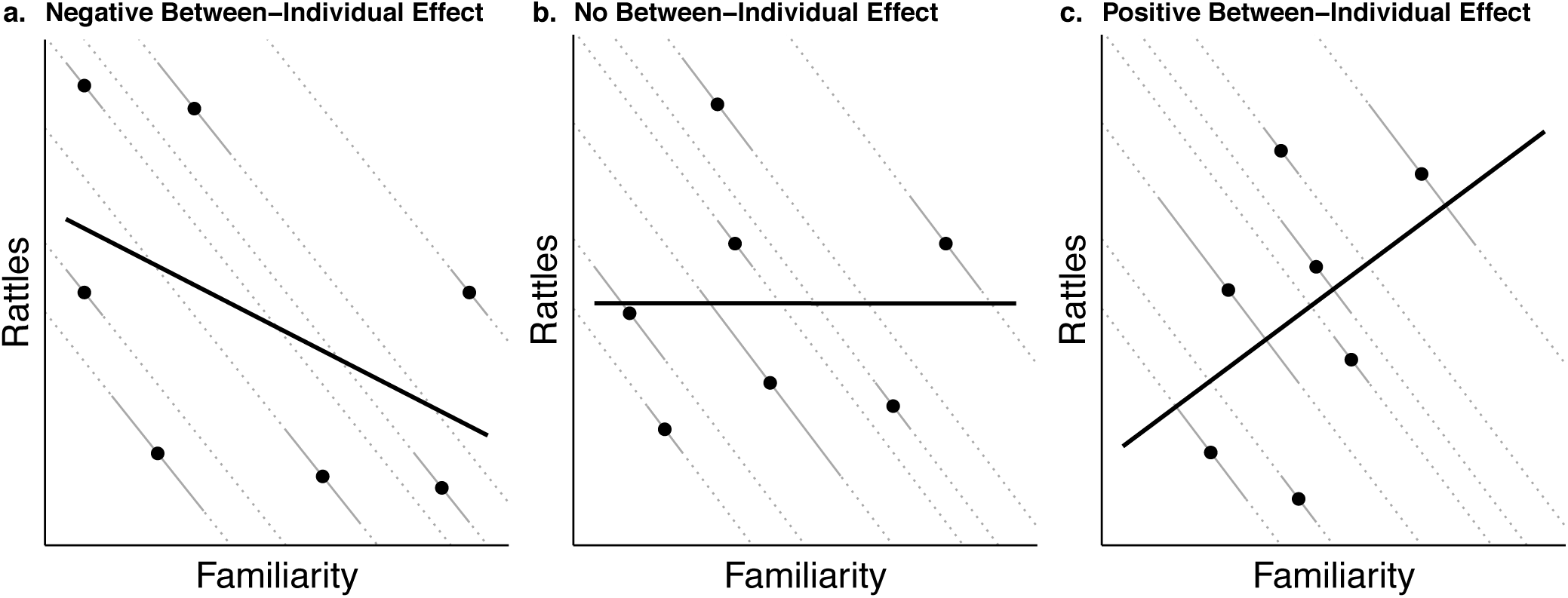
Three different scenarios for how variation in mean rattling rate (random intercepts) in combination with variation in data sampling structure might change our ability to detect among-individual effects when individuals have the same slope. We schematically depict the within-individual slopes (solid grey lines) of seven subjects (*j* = 1 to *j* = 7). The solid grey lines indicate the range over which each individual was sampled. Dotted lines provide an extension of these slopes to the edge of the figure. The among-subject slope (solid black line) is based on the association between *x̄_j_* and *ȳ_j_* as denoted by the filled black circles.

Although we have provided an explanation for the differences in our long-term and cross-sectional findings, there are a couple reasons why it is important to interpret our results with caution. First, there is potential for changes in rattling rates to be driven by effects of age rather than familiarity if the strength of territory defence declines with physical deterioration. We have done our best to account for this possibility in our analyses, but given that these variables are strongly correlated an experimental approach would prove useful in disentangling these effects as they are difficult to tease apart statistically. Second, it is worth addressing our use of multiple univariate analyses to test a single overarching hypothesis. Previous research in this study system has detected effects of the social environment on vigilance and nest use using a multivariate analysis (Dantzer et al., 2012), allowing us to make specific predictions about how squirrels should adjust patterns of nest use and vigilance in response to neighbourhood familiarity. Given this, we felt that analyzing the effects of familiarity on each of these behaviours individually provided a more elegant test of our hypothesis. However, it is important to be aware that our use of univariate analyses increases our chances of committing a Type I error by attributing variance as unique to a single response variable when it may in fact be shared (Huberty & Morris, 1989).

Despite these limitations, we believe that the results from our study, in particular the data for which we can estimate within-individual changes in behaviour, provide evidence that red squirrels are socially plastic. Furthermore, although we have not directly tested the fitness consequences of social plasticity, red squirrels reduced rattling rates, thereby spending less time on territory defence and potentially minimizing risk of detection by predators, under social conditions where intrusion risk was low (Siracusa, Boutin, et al., 2017a). This suggests that ‘asocial’ species can not only be socially responsive but also socially competent in their behaviour (Taborsky & Oliveira, 2013; 2012). Familiarity with conspecifics has long been assumed to be beneficial by allowing individuals to minimize energy expended on territory contests and increase time devoted to reproduction and growth (Getty, 1987; Temeles, 1994; Ydenberg et al., 1988). Evidence for reduced aggression toward familiar conspecifics is taxonomically widespread (reviewed in Temeles, 1994). However, these studies have typically been focused on documenting behavioural changes on short timescales through exposure to an experimental stimulus. Our study is one of few to demonstrate that natural variation in neighbourhood familiarity has direct consequences for behavioural time budgets by allowing individuals with familiar neighbours to reduce territory defence and increase time spent in nest. A handful of previous studies have demonstrated similar patterns in wild populations under natural social conditions. Willow ptarmigan (*Lagopus lagopus*) males were found to spend significantly more time engaged in territorial border disputes when they had more new neighbours (Eason & Hannon, 1994). In Seychelles warblers (*Acrocephalus sechellensis*), living near familiar individuals provided important benefits by reducing immediate energetic costs through fewer physical fights (Bebbington et al., 2017).

Additionally, recent research has increasingly noted the importance of group composition in shaping individual behaviour (Farine, Montiglio, & Spiegel, 2015). For example, nutmeg mannikins (*Lonchura punctulata*) have been shown to forego consistent individual differences in scrounger-forager tactics when flock composition changes, and to adjust their social strategy according to frequency-dependent pay-offs (Morand-Ferron, Wu, & Giraldeau, 2011). Water striders (*Aquarius remigis*) also show plasticity in aggression and activity in response to the presence of hyperaggressive individuals in the group (Sih et al., 2014) or changes in male-male competition (Montiglio et al., 2017). Although territorial species do not act in clearly defined, discrete units, we have demonstrated that red squirrels show similar social plasticity in response to the composition of neighbouring territory holders at the scale of the acoustic social environment (i.e. 130 m radius). Our results emphasize that the composition of neighbouring conspecifics, in addition to quantity of individuals in the social environment (Dantzer et al., 2012), can shape the behaviour of territorial species.

## Conclusion

The advantage of territoriality is contingent on the benefits of resource acquisition outweighing the costs of defending those resources from competitors (Brown, 1964; Schoener, 1987). Although the dear-enemy effect has been a well-recognized phenomenon in territorial species for several decades (Temeles, 1994), the relative importance of social interactions for maintaining this cost-benefit ratio under natural conditions has rarely been explored. Here we show that red squirrels adjust their behaviour in response to the familiarity of their social environment and that squirrels living in unfamiliar social neighbourhoods pay a cost in time by increasing rattling rates three-fold and reducing nest use by approximately 25% in order maintain their territory under conditions of high intrusion risk. Importantly, our results suggest that behavioural plasticity in this species cannot be estimated by comparing differences in behaviour among-individuals, emphasizing the need to have observations from the same individuals across multiple social environments in order to detect these behavioural patterns.

Taken altogether, these results provide evidence that territorial species have the capacity to assess and respond to nuanced changes in their social environment, despite not typically being considered to engage in important social interactions. Social relationships may therefore be more important than previously appreciated for apparently ‘asocial’ species. Several studies have demonstrated that familiarity with territory neighbours can have fitness benefits, including increased growth rate and survival (Höjesjö, Johnsson, Petersson, & Järvi, 1998; Seppä, Laurila, Peuhkuri, Piironen, & Lower, 2001), reduced telomere attrition (Bebbington et al., 2017) and increased reproductive success (Beletsky & Orians, 1989; Grabowska-Zhang et al., 2012). Given that these long-term social relationships have been demonstrated to affect both intrusion risk (Siracusa, Boutin, et al., 2017a) and behavioural time budgets (this study) in red squirrels, it seems likely that familiarity with territory neighbours could also have important fitness consequences for this species, but this has yet to be explored.

## Acknowledgments

We thank all of the field technicians who have contributed to the long-term KRSP database over the years. We are particularly indebted to M.Thorpe, Y.Sun and J. Robertson for their endless hours of squirrel watching and for waking up at unreasonable hours of the morning to deploy audio recorders. We acknowledge that this study was conducted on Champagne and Aishihik First Nations land and thank Agnes MacDonald and her family for long-term access to her trapline. This research was supported by funding from the Natural Sciences and Engineering Research Council of Canada (A.G. McAdam; RGPIN-2015-04707), as well as Grants-in-Aid of research from the American Society of Mammalogists and the Arctic Institute of North America (E.R. Siracusa). This is publication number XX of the Kluane Red Squirrel Project.

